# Growth media affects susceptibility of air-lifted human nasal epithelial cell cultures to SARS-CoV2, but not Influenza A, virus infection

**DOI:** 10.1101/2023.07.31.551381

**Authors:** Jessica D. Resnick, Jo L. Wilson, Eddy Anaya, Abigail Conte, Maggie Li, William Zhong, Michael A. Beer, Andrew Pekosz

## Abstract

Primary differentiated human epithelial cell cultures have been widely used by researchers to study viral fitness and virus-host interactions, especially during the COVID19 pandemic. These cultures recapitulate important characteristics of the respiratory epithelium such as diverse cell type composition, polarization, and innate immune responses. However, standardization and validation of these cultures remains an open issue. In this study, two different expansion medias were evaluated and the impact on the resulting differentiated culture was determined. Use of both Airway and Ex Plus media types resulted in high quality, consistent cultures that were able to be used for these studies. Upon histological evaluation, Airway-grown cultures were more organized and had a higher proportion of basal progenitor cells while Ex Plus-grown cultures had a higher proportion terminally differentiated cell types. In addition to having different cell type proportions and organization, the two different growth medias led to cultures with altered susceptibility to infection with SARS-CoV-2 but not Influenza A virus. RNAseq comparing cultures grown in different growth medias prior to differentiation uncovered a high degree of differentially expressed genes in cultures from the same donor. RNAseq on differentiated cultures showed less variation between growth medias but alterations in pathways that control the expression of human transmembrane proteases including *TMPRSS11* and *TMPRSS2* were documented. Enhanced susceptibility to SARS-CoV-2 cannot be explained by altered cell type proportions alone, rather serine protease cofactor expression also contributes to the enhanced replication of SARS-CoV-2 as inhibition with camostat affected replication of an early SARS-CoV-2 variant and a Delta, but not Omicron, variant showed difference in replication efficiency between culture types. Therefore, it is important for the research community to standardize cell culture protocols particularly when characterizing novel viruses.

## INTRODUCTION

Primary differentiated respiratory epithelial cell cultures have been widely used by researchers to study viral fitness and virus-host interactions, especially during the COVID19 pandemic ^1–8^. These cultures recapitulate important characteristics of the respiratory epithelium such as diverse cell type composition, polarization, and innate immune responses while maintaining desirable *in vitro* characteristics such as being relatively quick and easy to grow^1–8^. While immortalized cell cultures can be useful for studying molecular virological phenotypes, differentiated primary cell cultures are preferred for investigations of host-virus interactions, receptor usage, cell tropism, and innate responses ^9^. However, standardization and validation of these cultures remains an open issue ^10,11^.

The upper respiratory tract is made up of five major epithelial cell types-basal, suprabasal, club, goblet, and ciliated ^12^. These cell types represent a continuum of differentiation states and proportions of each vary throughout the respiratory tract ^9,12^. The cell tropism of respiratory viruses can vary across the respiratory tract and is usually based on expression of their preferred entry factors ^9,13^. For example, Influenza A viruses (IAV) which use sialic acid glycan receptors predominantly target ciliated cells where these are most highly expressed, while the most susceptible cell types to SARS-CoV-2 (SCV2) virus infection are ciliated and goblet cells which are not necessarily the highest expressors of the ACE2 receptor SCV2 uses for entry ^14,15^.

The most commonly used media for establishing cultures at the air-liquid interface is BEGM, but a more recently available media, Pneumacult, is gaining popularity due to the fact that it promotes development of goblet cells ^12^. Precise components and concentrations of commercial media are not available to most investigators, necessitating direct comparisons of cultures that have been propagated and differentiated using different media and growth conditions. Previous work has shown that differentiation media influences final culture morphology and cellular responses to viral infection but has no impact on infectious virus production or ciliation phenotypes ^16–18^. Other studies have shown that expansion media can impact the number of successful passages of progenitor cells ^19^.

Due to the impact of the ongoing COVID19 pandemic on the availability of patient-derived airway epithelial cultures, our laboratory turned to commercial sources of respiratory epithelial cells. In this study, we compared two different ways of expanding these cells prior to differentiation and measured the resulting impact on final culture organization, cell type proportions, and response to respiratory virus infection. We find that the effects of growth media persist through the differentiation process, leading to culture differences that contribute to differential susceptibility to SCV2, but not IAV, infection, likely through expression of key entry cofactors. This work highlights the importance of independent comparisons of cell culture reagents, and the need for standardization between studies when using this data to inform public health decisions.

## METHODS

### Cell Culture

Vero-E6 over expressing TMPRSS2 cells (VT; Japanese Collection of Research Bioresources Cell Bank, JCRB1819) ^20^. VT cells were cultured in Dulbecco’s Modified Eagle Medium (DMEM, Gibco) with 10% fetal bovine serum (FBS, Gibco Life Technologies), 100 U penicillin/mL with 100 µg streptomycin/mL (Quality Biological), 2 mM L-Glutamine (Gibco Life Technologies), and 1mM Sodium Pyruvate (Sigma) at 37°C with air supplemented with 5% CO2. Infectious medium specific for SCV2 (IM-SCV2) was used in all infections and consists of DMEM with 2.5% FBS, 100U penicillin/mL with 100 µg streptomycin/mL, 2 mM L-Glutamine, and 1 mM Sodium Pyruvate.

Madin-Darby canine kidney (MDCK) cells were cultured in Dulbecco’s Modified Eagle Medium (DMEM, Sigma-Aldrich) with 10% fetal bovine serum (FBS, Gibco Life Technologies), 100U penicillin/mL with 100 μg streptomycin/mL (Quality Biological), and 2 mM L-Glutamine (Gibco Life Technologies) at 37 °C with air supplemented with 5% CO_2_. Infectious medium for IAV (IM-IAV) was used in all infections and consists of DMEM with 4 µg/mL N-acetyl trypsin (NAT), 100 u/ml penicillin with 100 µg/ml streptomycin, 2 mM L-Glutamine and 0.5% bovine serum albumin (BSA) (Sigma).

Human nasal epithelial cells (hNEC) (Promocell, lot 466Z007, 466Z004, and 453Z019) were grown to confluence in 24-well Falcon filter inserts (0.4-µM pore; 0.33 cm^2^; Becton Dickinson) using PneumaCult™-Ex Plus Medium (Stemcell) or the Airway Epithelial Cell Grow Medium Kit (Promocell). Hereafter, the two medias will be referred to as Ex Plus and Airway media respectively. Donor 466Z007 was a 48-year-old Caucasian male, never smoker, and SARS-CoV-2 negative one day before collection. Donor 453Z019 was a 32-year-old Caucasian male. Donor 466Z004 was a 43-year-old Caucasian male. Confluence was determined by a transepithelial electrical resistance (TEER) reading above 250Ω by Ohm’s law method ^21^ and by examination using light microscopy and a 10x objective. The cells were then differentiated at an air-liquid interface (ALI) before infection, using ALI medium as basolateral medium as previously described ^1,10^. Briefly, both apical and basolateral media were removed and ALI differentiation media (Stem Cell Technologies, Pneumacult ALI Basal Medium) supplemented with 1X ALI Maintenance Supplement (StemCell Technologies), 0.48 µg/mL Hydrocortisone solution (StemCell Technologies), and 4 µg/mL Heparin sodium salt in PBS (StemCell Technologies) was replaced on the basolateral side only. Fresh media was given every 48 hours. Hereafter, differentiation media will be referred to as ALI media. Once mucus was visible, apical washes were performed weekly with PBS to remove excess mucus. Cells were considered fully differentiated after 3 weeks and when cilia were visible using light microscopy and 10x objective. All cells were maintained at 37°C in a humidified incubator supplemented with 5% CO2.

### Virus Seed Stock and Working Stock Generation

The SARS-CoV-2 virus used in this study, designated SARS-CoV-2/USA/ DC-HP00080/2020 (B.1; GISAID EPI_ISL_438237), was isolated from samples obtained through the Johns Hopkins Hospital network ^22^. For virus working stocks, VT cells in a T75 or T150 flask were infected at an MOI of 0.001 with virus diluted in IM. After a one-hour incubation at 33 °C, the inoculum was removed and IM was added (10 ml for T75 and 20 ml for a T150 flask). When cytopathic effect was seen in approximately 75% of the cells, the supernatant was harvested, clarified by centrifugation at 400 g for 10 minutes, aliquoted and stored at −65C. Delta B.1.617.2 (AY.106) (SARS-CoV2/USA/MD-HP05660/2021; GISAID EPI_ISL_2331507) and Omicron B1.1.529 (BA.1) (hCoV19/USA/MD-HP20874/2022; GISAID EPI_ISL_7160424) viruses used were generated in the same manner.

The Influenza A Virus used was A/Baltimore/R0243/2018 (H3N2 clade 3C.3a) (GISAID EPI_ISL_17034889) was also isolated from samples obtained through the Johns Hopkins Hospital network as part of the CEIRS network ^23^. For virus working stocks, MDCK cells in a T150 flask were infected at an MOI of 0.001 with virus diluted in IM. After one hour, the inoculum was removed, and fresh IM was added. When cytopathic effect was seen in approximately 50% of cells, the supernatant was harvested, aliquoted, and stored at −65 °C.

### TCID_50_ Assay

VT or MDCK cells were grown to 90-100% confluence in 96-well plates. After being washed twice with PBS+, ten-fold serial dilutions of the viruses in IM were made and each dilution was added to 6 wells. The plates were incubated at 37 °C with 5% CO_2_ for 5 days. The cells were fixed by adding 75 µL of 4% formaldehyde in PBS per well overnight and then stained with Napthol Blue Black solution overnight. Endpoint values were calculated by the Reed-Muench method ^24^.

### Low Multiplicity of Infection (MOI) infections

For hNEC infections, an MOI of 0.1 and 1 TCID50 per cell was used for IAV and SCV2 respectively. The basolateral media was collected, stored at −65 °C, and replaced with fresh media every 48 hours. The apical side of the transwell was washed 3 times with IM, with a 10-minute incubation at 37 °C in between each. The virus inoculum was diluted in its matched virus IM (mock used SCV2 IM) and 100 µL was added to the apical side of cells and allowed to incubate for 2 hours. The inoculum was then removed, the cells washed 3 times with PBS-, and returned to the incubator. At 48 hours post infection, a 10-minute apical wash was performed with IM and collected and stored at −65 °C. Infectious virus particle production in apical washes was quantified using TCID50 on VT or MDCK cells for SARS-CoV2 and Influenza A Viruses respectively.

### Cytokine Secretion

Secreted interferons, cytokines, and chemokines were quantified from the basolateral samples at 0 and 48 hours post infection from the hNEC infections. Measurements were performed using the V-Plex Human Chemokine Panel 1 (CCL2, CCL3, CCL4, CCL11, CCL17, CCL22, CCL26, CXCL10, and IL-8) (Meso Scale Discovery) and the DIY Human IFN Lambda 1/2/3 (IL-29/28A/28B) ELISA (PBL Assay Science) according to the manufacturer’s instructions. Each sample was analyzed in duplicate. Heatmaps were generated and hierarchical clustering was performed using the R package “pheatmap”.

### Imaging

The hNEC cultures were infected with SARS-CoV-2 and IAV at an MOI of 1 and 0.1 respectively. At 48 hours post infection, the wells were washed twice with PBS- and then fixed using 4% paraformaldehyde in PBS on both the apical and basolateral sides for 20 minutes at room temperature. The wells were then washed twice with PBS- and stored at 4 °C in PBS-until ready to be stained.

hNEC wells were then permeabilized and blocked with PBS containing 0.5% Triton X-100 and 5% BSA. The samples were incubated with 2.25 µg/ml anti-TMPRSS2 (Proteintech, Cat# 14437-1-AP), anti-ACE2 1 µg/ml (Genetex, Cat# GTX101395), anti-TMPRSS11E protein 9 µg/ml (Invitrogen, PA5-50809), and anti-β-Tubulin IV 5 µg/ml (Novus, Cat# NBP2-00812), 1.65 µg/ml SCV2 (GTX135357), or 2.15 µg/ml IAV (GTX125989) primary antibodies overnight at 4 °C. Fluorescently labeled secondary antibodies AF488 (4 µg/ml) (ThermoFisher, A11013) and AF647 (4 µg/ml)(ThermoFisher, A21235) were used as secondary stains for 1 hour at room temperature. After washing, hNECs were incubated with Hoechst 33258 (2 µg/ml) (Invitrogen, H3569) and Rhodamine Phalloidin (100 nM)(Cytoskeleton, #PHDR1) for 30 minutes at room temperature. The slides were sealed with a coverslip using Prolong glass antifade medium (Invitrogen, P36984). Images were acquired using a Zeiss LSM700 at 63x magnification with 1 µm z-stack sections. Mean fluorescence intensity per section was quantified using ImageJ.

Individual transwells containing hNEC were submerged in 10% Neutral Buffered Formalin (Leica 3800598) for 30 mins and went through a series of dehydration processes in 70%-100% ethanol (Fisher BioReagents BP2818500), and xylenes (Fisher Chemical X5-500). The dehydrated hNEC transwell membrane was then separated using a surgical blade and incubated in 65 °C paraffin (Leica EM-400 3801320) for 30 mins. Samples were then embedded and sectioned at 4.5 µM (Leica HistoCore 149AUTO00C1) and transferred to a 42 °C distilled water bath and collected using positively charged slides. Sections later were processed using routine H&E staining (Vector Laboratories H3502) and cover slipped for imaging. Images were obtained using EVOS XL Core microscope at 20X magnification.

### Flow Cytometry

Fully differentiated hNECs with either differentiation condition (Airway or Ex-Plus Media) were harvested from the apical membrane into a single cell suspension with a 30-minute incubation in 1X TrypLE (Gibco 12563011). After cells are trypsinized and resuspended in a trypsin stop solution (10% FBS in PBS, Thermofisher, Gibco, Lot:2193952RP). The cells were then washed three times in 1X PBS and resuspended in 1 mL PBS (centrifuge at 2500 RPM between wash steps). Appropriate control and sample tubes were then stained with AQUA viability dye (Invitrogen L34965) 1 µL/1×10^6^ cells for 30 minutes at room temperature. Cells were then washed and resuspended in BD Fixation/Permeabilization solution (BD Biosciences 554714) and incubated for at least 30 minutes at 4 °C. Cells were washed with BD Perm/Wash Buffer x2 and centrifuged at 2500 RPM at 4 °C for 5 minutes. Cells were then resuspended in BD Perm/Wash Buffer with 7% NGS (Sigma Aldrich G9023) and incubated for 1 hour at 4 °C. Cells were washed with BD Perm/Wash Buffer x2 and centrifuged at 2500 RPM at 4 °C for 5 minutes. Appropriate sample tubes were incubated with primary antibodies for one hour at room temperature. Antibodies are diluted into BD Perm/Wash buffer at appropriate concentrations. Final staining volume is 200 µL. Cells were washed with BD Perm/Wash Buffer x2 at 2500 RPM and centrifuged at 4 °C for 5 minutes. Appropriate sample tubes were incubated with secondary antibodies for 30 minutes at room temperature. Cells were washed with BD Perm/Wash Buffer x2 at 2500 RPM and centrifuged at 4 °C for 5 minutes. Appropriate sample tubes were incubated with conjugated antibodies for 30 minutes at room temperature. Cells were washed with BD Perm/Wash Buffer x2 and centrifuged at 2500 RPM at 4 °C for 5 minutes. Cells were resuspended in FACS Buffer (0.3% BSA in 1X PBS, BSA: Sigma Aldrich A9418, PBS PH 7.4: Gibco 10010072) and filtered through a 35 µM strainer cap into FACS tubes just prior to the run. Cell suspensions were run on a BD LSRII Flow Cytometer using DIVA software. Single stained cells were used as controls and fluorescence minus one controls were used to assist in gating. Data analysis was completed on FlowJo V10. Gating strategy employed was as follows: exclusion of debris, single cells, and Aqua – cells (LIVE cells) (supp Fig 1).

#### Antibody List

**Table 1:** Antibodies used for Flow cytometry.

|  |  |  |  |
| --- | --- | --- | --- |
| Instrument: BD LSRII |  |  |  |
| Antibody/Probe/Clone | Fluorophore | Catalog # | Staining Concentration |
| Anti-Acetylated α- | AF647 | Santa-Cruz<br>Biotechnology SC- | 200 ng/mL |
| Tubulin |  | 23950 |  |
| Rabbit anti-ACE2 | Primary | Proteintech Cat. No 21115-1-AP | 0.5 µg/mL |
| Rabbit anti-TMPRSS2 | Primary | Proteintech Cat. No 14437-I-AP | 0.5 µg/mL |
| Mouse IgG 1 Anti-TMPRSS11E Clone TM191 | Primary | 392002 | 0.01 mg/mL |
| MUC5AC Monoclonal Antibody Clone: 45M1 | Primary | Invitrogen MA5-12178 | 2 µg/mL |
| CD271 (NGF Receptor) Monoclonal Antibody (ME20.4), PE, eBioscience | Conjugated-PE | Invitrogen 12-9400-42 | 0.5 µg/mL |
| Goat Anti-Mouse (Secondary Ab for MUC5AC probe) Clone: Poly4503 | BV605 | Biolegend 405327 | 0.2 µg/mL |
| Mouse Anti-Beta Tubulin-IV | AF488 | Novus Bio NBP2-74713AF488 | 0.78 µg/mL |
| Goat anti-Rabbit IgG (H+L) Highly Cross-Adsorbed Secondary Antibody | AF488 | Catalog # A-11008 | 1 µg/mL |
| Goat anti-Mouse IgG (H+L) Highly Cross-Adsorbed Secondary Antibody | AF488 | Catalog # A32723 | 1 µg/mL |
| Live/Dead Discriminator | AQUA | Invitrogen L34965 | 1 µL/10<sup>6</sup> cells |

### RNA-Sequencing

Total RNA was extracted and purified from hNECs using Trizol reagent and the PureLink RNA Mini kit, including on-column DNAse treatment (Invitrogen/ThermoFisher). Quantitation of Total RNA was performed with the Qubit BR RNA Assay kit and Qubit Flex Fluorometer (Invitrogen/ThermoFisher), and quality assessment performed by RNA ScreenTape analysis on an Agilent TapeStation 2200. Unique Dual-index Barcoded libraries for RNA-Seq were prepared from 100 ng Total RNA using the Universal Plus Total RNA-Seq with NuQuant Library kit (Tecan Genomics), according to manufacturer’s recommended protocol. Library amplification was performed for 16 cycles, as optimized by qPCR.

Quality of libraries was assessed by High Sensitivity DNA Lab Chips on an Agilent BioAnalyzer 2100. Quantitation was performed with NuQuant reagent, and confirmed by Qubit High Sensitivity DNA assay, on Qubit 4 and Qubit Flex Fluorometers (Invitrogen/ThermoFisher). Libraries were diluted, and equimolar pools prepared, according to manufacturer’s protocol for appropriate sequencer. An Illumina iSeq Sequencer with iSeq100 i1 reagent V2 300 cycle kit was used for final quality assessment of the library pool. For deep RNA sequencing, a 200 cycle (2×100bp) Illumina NovaSeq S2 run was performed at Johns Hopkins Genomics, Genetic Resources Core Facility, RRID:SCR_018669.

iSeq and NovaSeq data files were uploaded to the Partek Server and analysis with Partek Flow NGS software, with RNA Toolkit, was performed as follows: pre-alignment QA/QC and trimming of reads. Following this, sequences were uploaded to the Beer lab cluster for further analysis^25^.

Sequences were first checked for quality using FastQC ^26^. All sequences were determined to be of good quality and were then aligned using HISAT2 to the GRCH38 genome ^27^. SAM files were then converted to BAM using samtools ^28^. A gene-count matrix was then generated from BAM files using featureCounts, and differential expression analysis was performed using DESeq2 in R ^29,30^. Pathway analysis of differentially expressed genes was also performed using clusterProfiler and gProfiler ^31,32^. For detailed methods and a full list of packages used please see https://github.com/JRes9/Resnicketal_Media_2023 (Accessed on July 24, 2023).

All sequence files and sample information are available at NCBI Sequence Read Archive, NCBI BioProject: PRJNA946012.

### RNA extraction and qPCR

RNA was extracted from hNECs using Trizol (Invitrogen, 15596026) and the PureLink RNA Mini Kit with on column DNase treatment (Invitrogen, 12183018A) according to manufacturer protocol. RNA was then converted to cDNA using the High-Capacity cDNA Reverse Transcription Kit (ThermoFisher, 4368814) according to manufacturer protocol. cDNA was diluted 1:10. qPCR was then run using Taqman reagents according to manufacturer protocol (Master Mix: applied biosystems, 4369016). Probes used were as follows: TMPRSS2 (applied biosystems, Hs00237175_m1), TMPRSS11E (applied biosystems, Hs01070171_m1), ACE2 (applied biosystems, Hs01085333_m1), and GAPDH (applied biosystems, hs02786624_g1).

### Drug Inhibition Assays

The drugs used for inhibition were as follows:

Aloxistatin (E64D, cathepsin inhibitor)-25 mg from MedChem Express (CAT HY-100229/CS-5996, LOT 114325), Molecular weight 342.43

Camostat mesylate (TTSP inhibitor) −10 mg from SIGMA (CAT SML0057, Batch 0000114299), Molecular weight 494.52

The vehicle used for both drugs was DMSO. Drugs were maintained at −20 °C in both high concentration (10 mM) and low concentration (100 µM) stocks.

Fully differentiated hNEC wells were first treated with a range of drug concentrations for 72 hours to determine cytotoxicity. Fresh dilutions of each drug in media were made daily. Cell viability was measured using alamarBlue (ThermoFisher) according to manufacturer instructions. Briefly, for each timepoint alamarBlue was added to the basolateral media at 10% of the total volume, then incubated at 37 °C for 4 hours. The basolateral media replaced with fresh media containing drug, and the old media was then aliquoted into 3 wells of a 96 well plate and absorbance read in triplicate. Results were normalized to both an untreated well and a media only well as positive and negative controls respectively.

Once baseline viability was determined (supp Figure 2), cells were pretreated with the indicated concentrations of drug in the basolateral media for 24 hours prior to infection. Basolateral media was then replaced with fresh media containing drug and infection was performed as described above. Viability was determined by alamarBlue at 48 hours post infection after collection of the apical wash. Infectious virus production in apical wash was determined by TCID50.

## RESULTS

Matched lots of hNECs were grown to confluence on aTranswell in either Airway or Ex Plus media before being differentiated using ALI media. Approximately 21 days post establishment of the air-liquid interface (ALI), when cultures were producing mucus and had visible cilia under the microscope, wells were section and stained by H & E to observe cell type proportions and overall organization (Fig 1 A and B). Airway grown cultures had a more distinct basal cell layer, greater overall organization, and less terminally differentiated cells. Ex plus grown cultures had more cells overall (despite air lift occurring at equal confluence) and a greater proportion of terminally differentiated ciliated and mucus producing cells. Cell type proportions were also evaluated by flow cytometry (Fig 1C) which confirmed that while Airway grown cultures had a greater proportion of basal progenitor cells, Ex Plus grown cultures had a higher proportion of terminally differentiated cell types, particularly ciliated and goblet cells.

**Figure 1.**
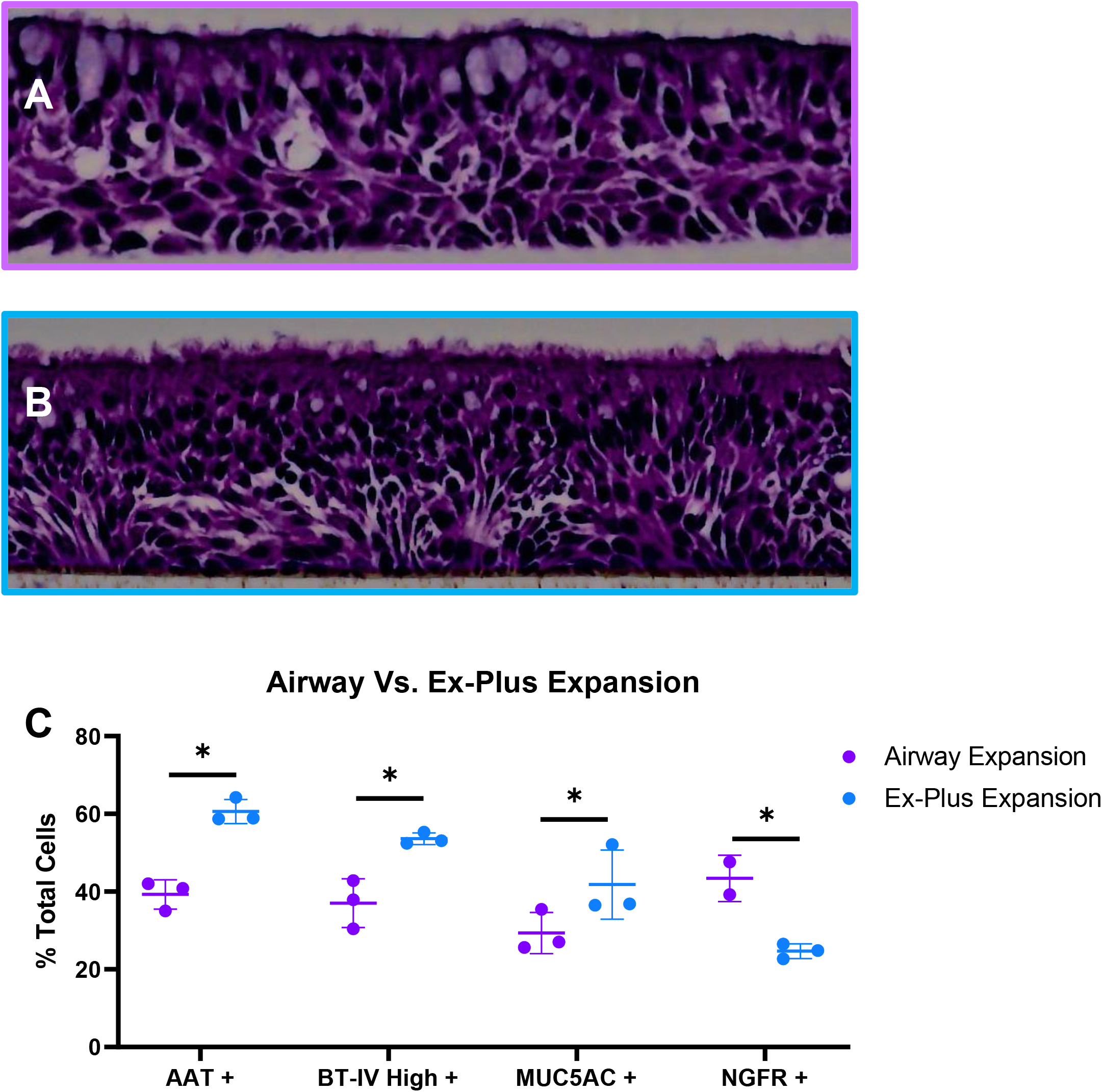
Cell organization and proportions present after differentiation. Fully differentiated human nasal epithelial cellcultures that had been expanded with either Airway (A) or Ex Plus (B) media were fixed, sectioned, and H & E stained. Cell-type proportions were determined in separate wells using flow cytometry (C). Cells were gated by excluding debris and single cells, then staining for the markers indicated. Percent of total cells staining with each marker was calculated to account for different cell numbers between conditions. Data is pooled from 3 wells of each condition, with each experiment performed two times. Data from one representative experiment is shown. *p<0.05 by two-way ANOVA with Tukey’s posttest.

To investigate whether growth media impacts susceptibility to infection, matched Airway and Ex Plus cultures were infected with a clinical isolate of either Influenza A (H3N2, IAV) or SARS-CoV-2 (B.1, SCV2) virus. At 48 hours post infection (HPI), there were no apparent differences seen in the number of IAV infected cells between cultures, but there were more SCV2 infected cells in Ex Plus grown cultures than Airway grown (Fig 2A). Additionally, Ex Plus grown cultures produced significantly more infectious SCV2 virus 48 HPI than Airway cultures but there was no difference in IAV production between the cultures (Fig 3).

**Figure 2.**
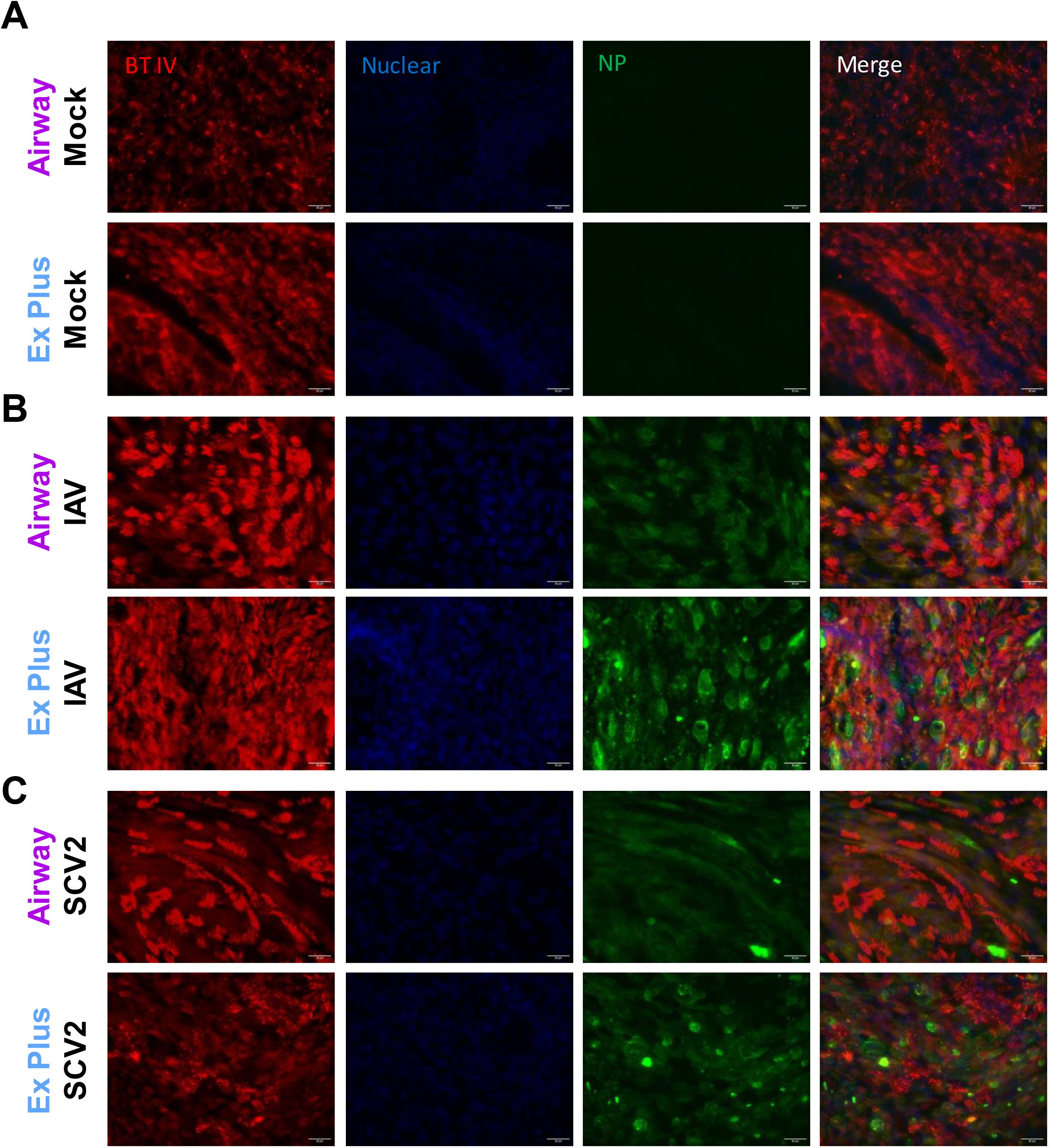
Infection of cultures with Influenza A Virus (IAV) or SARS-CoV-2 (SCV2). Airway or Ex Plus grown *c*ultures were either unifected (A) or infected with IAV (B) or SCV2 (C) at an MOI of 0.5 for 48 hours before being fixed and stained for cell and viral markers. Three wells per condition were used in each experiment and the experiment was repeated once. Representative data is shown.

**Figure 3.**
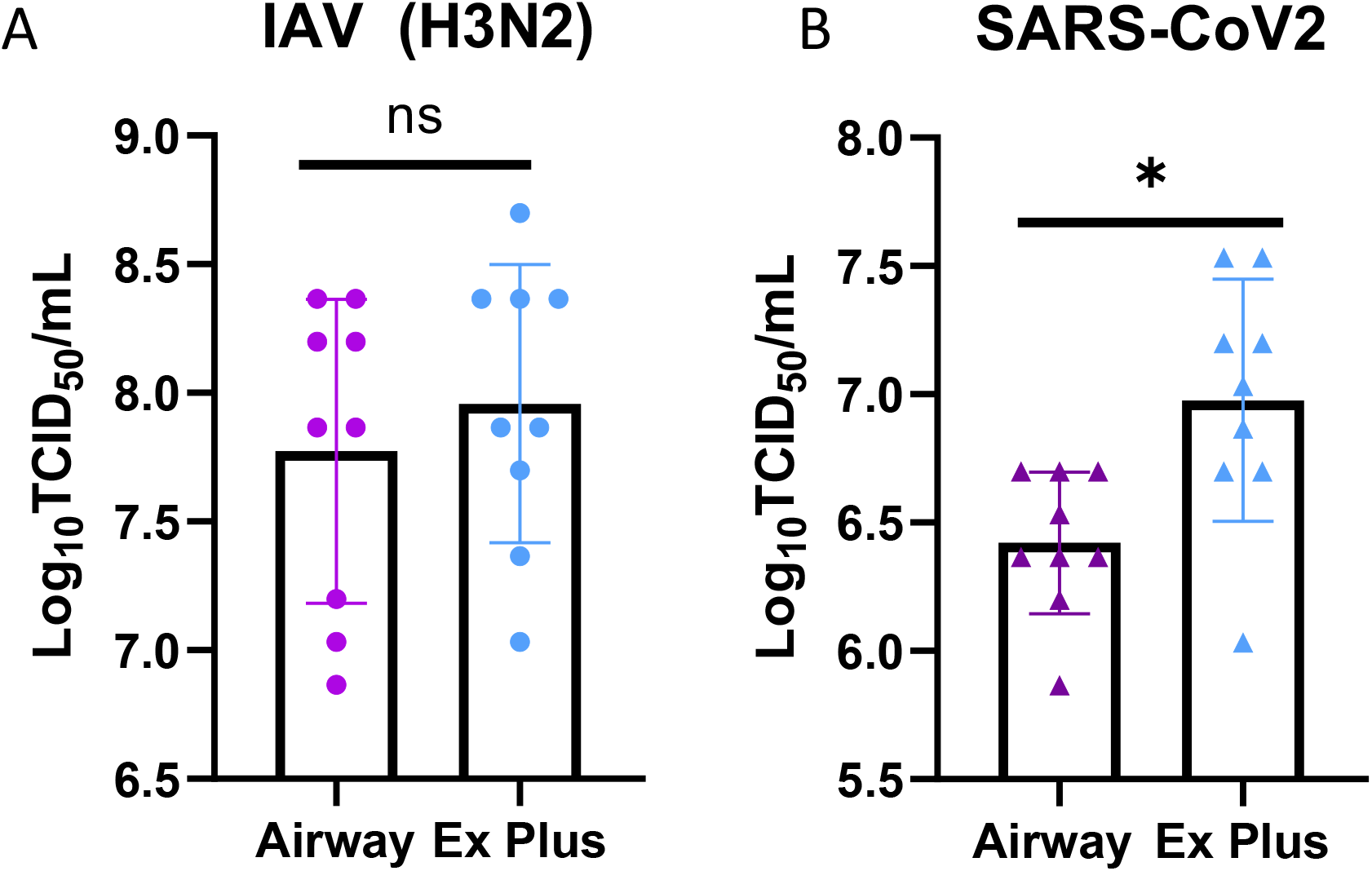
Infection of cultures with Influenza A Virus (IAV) or SARS-CoV-2 (SCV2). Cultures were infected with either IAV (A) or SCV2 (B) at an MOI of 0.1 or 1.0, respectively. Apical washes were collected, and infectious virus was determined by TCID50 at 48 hpi. Data are pooled from three independent experiments each with n=3 wells per virus (total n=9 wells per virus). *p<0.05 one-way ANOVA with Bonferroni correction.

To determine whether growth media was altering virus infection induced cytokine production, basolateral supernatant was collected from mock infected, IAV infected, and SCV2 infected cultures and a panel of pro-inflammatory cytokines, chemokines and interferon lambda production was measured (Fig 4) ^33,34^. Samples appear to cluster by treatment, rather than growth media, suggesting that induced cytokine and chemokine patterns are not significantly altered by growth media.

**Figure 4.**
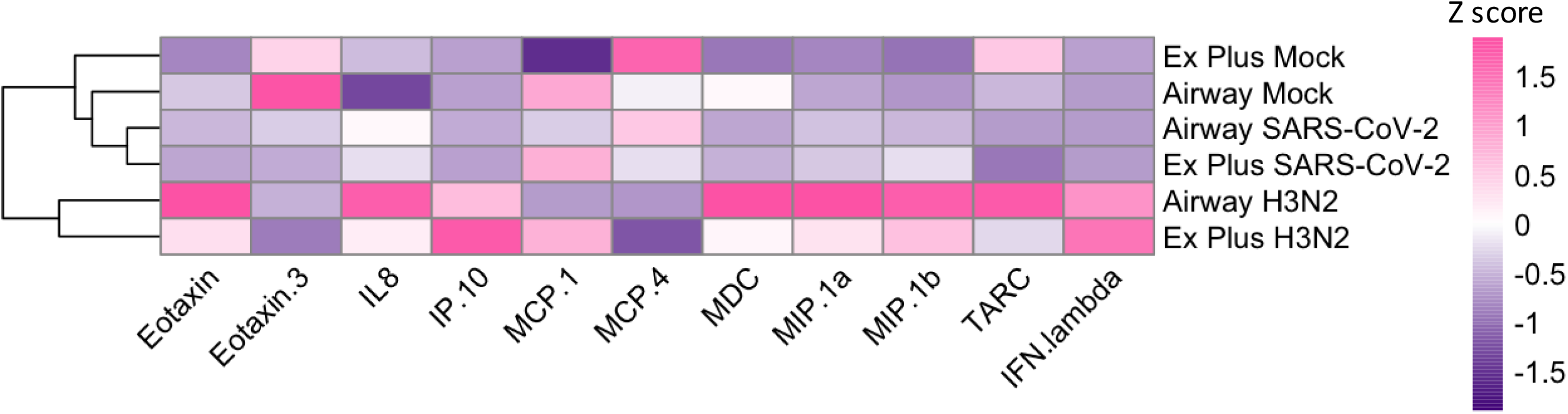
Basal and induced cytokine production in cultures. Basolateral secretions of cytokines, chemokines, and interferon lambda were measured 0 and 48HPI during infections with either IAV or SCV2 (n=3 wells per replicate, 9 wells total). Values were averaged and then scaled to calculate z-score. Hierarchical clustering was performed based on sample.

RNA-sequencing was then performed to identify expression differences between cultures from the same donor that were propagated in different growth media. Both Airway and Ex Plus grown cultures were maintained in the same ALI media for 3 weeks prior to collection, so differences were expected to be minimal and highly impactful. Cultures were collected on the last day of growth media (day 10-12) and when fully differentiated (~3 weeks post air lift). Differential expression analysis between fully differentiated cultures revealed that Ex Plus grown cultures had higher expression of TMPRSS11E than Airway grown, a serine protease that can prime the SCV2 spike protein and is most highly expressed in the upper airway (Figure 5) ^35^. Differential cofactor expression was confirmed using qPCR (supp Fig 3). Additionally, Airway grown cells showed increased expression of Pax6, which has been shown to negatively regulate TMPRSS2 in eye cells ^36^. Ex plus cells also have upregulated Six3, which regulates Pax6 ^37^. Taken together, these data indicated that SCV2 cofactor expression can be altered by the growth media used in the propagation phase of culturing.

**Figure 5.**
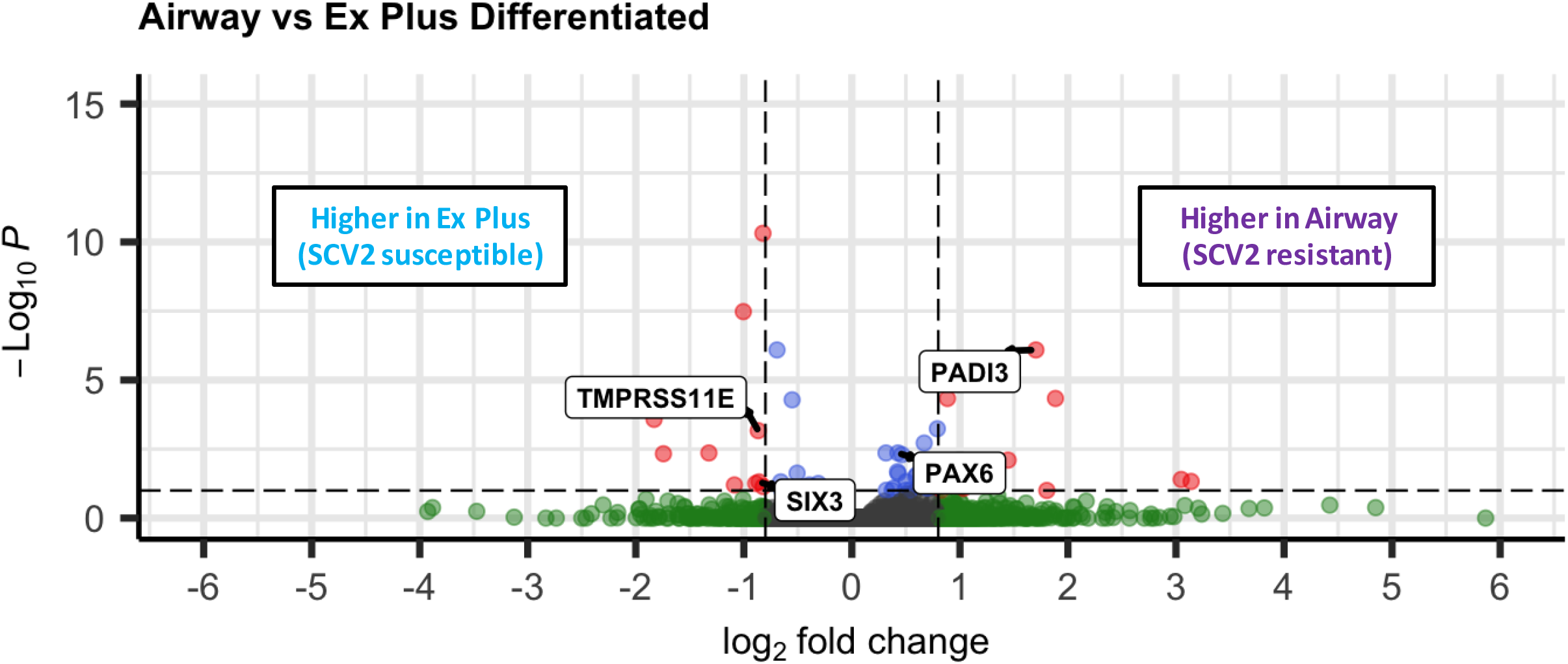
Differentially expressed genes between fully differentiated Ex Plus and Airway expanded cultures. Fully differentiated cultures were collected for bulk RNA-sequencing at day 21 post ALI. Data were pooled from three replicate wells. Log 2 fold change indicates the mean expression for a gene. Each dot represents one gene. Black dots indicate no significantly differential expression between Airway and Ex Plus expanded cultures. Blue dots indicate an adjusted p value <0.05. Green dots indicate an absolute log 2 fold change higher than 0.8. Red dots indicate both a significant p value and log 2 fold change.

In cultures that were harvested before ALI differentiation, the two growth media showed vastly different patterns of differentially expressed genes (Supp. Fig. 4). Pathway analysis showed an upregulation of pathways involved in ciliate-related functions and abnormal pulmonary functions in Ex Plus cultures (Supp. Fig. 5A) while pathways involved in cell adhesion dominated cultures grown in Airway media (Supp. Fig. 5B). The downregulated pathways also differed markedly depending on growth factor (Supp. Fig. 5C and D). This data indicate that while growth media can lead to markedly different gene expression patterns, the differentiation at ALI tends to minimize most but not all transcriptional differences.

SCV2 can use two different routes of viral entry. The late cleavage pathway, predicted to mostly be used in immortalized cells like Vero E6, involves endocytosis followed by priming of the S protein in the endosome by cathepsins ^35,38,39^. In contrast, the early cleavage pathway, predicted to be favored in the respiratory tract, involves priming by membrane-associated serine proteases and direct membrane fusion leading to genome release ^35,38,39^. To test which pathway (and related cofactor) is being utilized in the hNEC cultures, hNEC cultures were pretreated with either a cathepsin (E64D) or serine protease (Camostat) inhibitor and then infected with either IAV or SCV2 (Figure 6) ^35^. Treatment with the serine protease, but not cathepsin, inhibitor significantly reduced infectious virus production during SCV2 infection in both cultures (Figure 6A). However, Airway grown cultures were more sensitive to lower concentrations of camostat and showed a more significant reduction in infectious virus production (2.14 fold change reduction at high concentrations) than Ex Plus grown cultures treated the same (1.42 fold change reduction), with many wells having undetectable infectious virus. Additionally, while we see a similar trend of IAV infectious virus production being reduced by serine protease inhibition, it is to a significantly smaller extent (~1.2 fold for both media types). This is likely due to the fact that while IAV utilizes serine proteases for cleavage and viral entry, it is far more promiscuous in utilizing trypsin-like proteases in respiratory epithelial cells ^40^.

**Figure 6.**
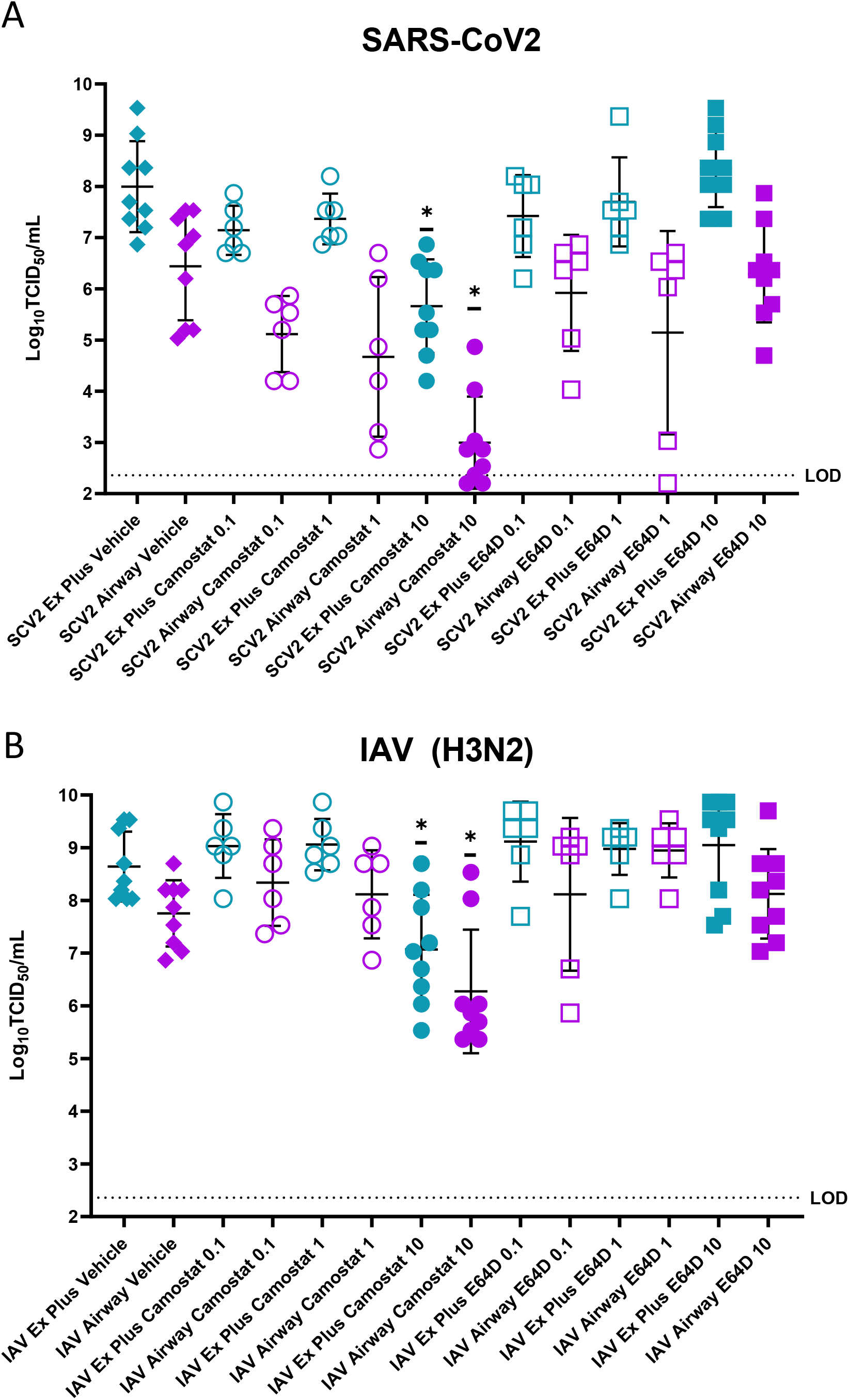
Protease inhibition during SCV2 or IAV infection of either Airway or Ex Plus expanded cultures. Cultures were pretreated with varying concentrations of either Camostat or E64D for 4 hours before being infected with the indicated virus. Apical washes were taken and infectious virus produced was quantified by TCID50 48 hours post infection. *p<0.05 (One-way ANOVA with Tukey’s posttest, compared to matched vehicle). Experiments were performed with n=3 replicates and the data from three experiments is shown.

Finally, different SARS-CoV-2 variants of concern have different entry pathway preferences ^41^. Delta variant viruses tend to use the early cleavage pathway, while omicron tend to use the late cleavage pathway ^41^. To further test that the early, but not late, cleavage pathway factors are impacted by growth media, airway or ex plus grown cultures were infected with either a parental, delta, or omicron variant virus and infectious virus production after 48 hours was determined. Infection with a parental or delta variant virus, predicted to prefer entry via the early cleavage pathway, produced more infectious virus in Ex plus grown cultures compared to airway grown (Fig 7). However, infection with omicron variant virus produced similar amounts of infectious virus in both culture types. These data again suggest that differences in serine protease expression or activity is likely driving differential susceptibility to and infectious virus production of SCV2 but not IAV virus.

**Figure 7.**
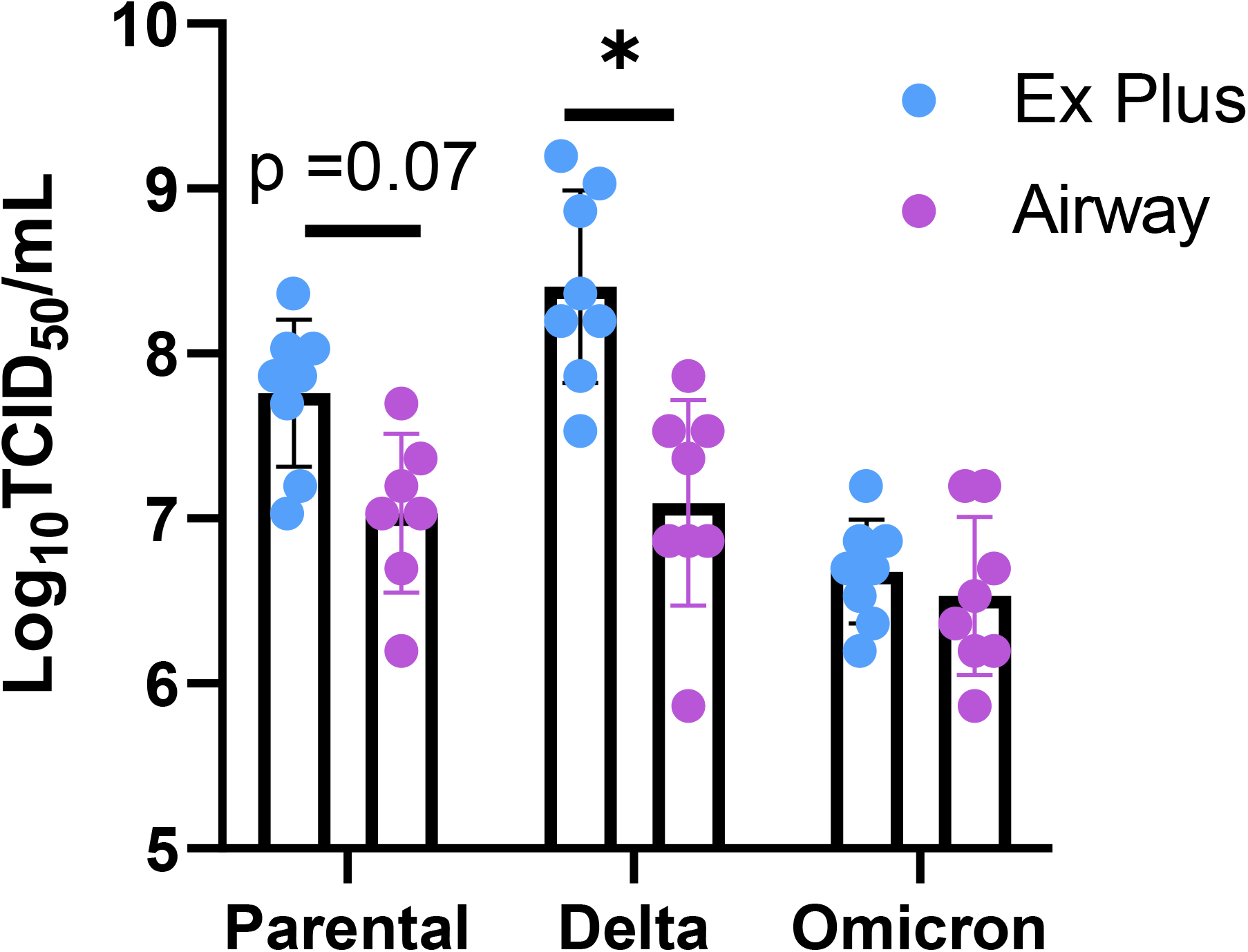
Comparison of susceptibility of Airway or Ex Plus expanded cultures to different SARS-CoV-2 variants of concern. Cultures were infected at an MOI of 0.1 with the indicated virus. Apical washes were taken 48 HPI and infectious virus was quantified by TCID50. Data are pooled from two independent experiments each with n=4 wells per virus (total n=8 wells per virus). *p<0.05 one-way ANOVA with Tukey’s posttest. Experiments were performed with n=3 replicates and the data from three experiments is shown.

## DISCUSSION

In this study, two different expansion medias were evaluated and the impact on the resulting differentiated culture was determined. Use of both Airway and Ex Plus media types resulted in high quality, consistent cultures that were able to be used for these studies. Upon histological evaluation, airway-grown cultures were more organized and had a higher proportion of basal progenitor cells while ex plus-grown cultures had a higher proportion of susceptible, terminally differentiated cell types. These differences may be a characteristic of different regions of the respiratory tract which should be taken into account during studies ^9,42,43^.

In addition to having different cell type proportions and organization, the two different growth medias led to cultures with altered susceptibility to infection with SCV2 but IAV. This cannot be explained by cell type proportion alone or we would expect that IAV would replicate more efficiently in Ex Plus grown cells which have a higher number of ciliated cells, its preferred host ^14^. RNA-seq analysis suggested that serine protease cofactor, rather than receptor or antiviral factor expression, led to this difference in replication of SCV2. Cofactor expression has previously been shown to impact viral entry pathway usage in immortalized cell types and is emerging as an important consideration during therapeutic development for COVID19 after the failure of hydroxychloroquine ^44–46^. Utilization of serine-protease cofactors in the hNEC culture system was confirmed using inhibitors of both cathepsins and serine proteases. Only treatment with the serine protease inhibitor impacted IAV or SCV2 replication, and it had a higher effect in the SCV2 infection setting. Furthermore, Airway grown cultures were more sensitive to lower concentrations of the serine protease inhibitor, again highlighting that there may be difference in either cofactor expression or activity between cultures. Finally, infection efficiencies of different SCV2 variants that have been previously shown to have different entry pathway preferences follow the predicted cofactor expression profiles of the different cultures ^41^. A representative delta variant, which have been shown to predominantly use the early cleavage pathway, replicate to significantly higher titers in Ex Plus grown compared to Airway grown cultures. In contrast, a representative omicron variant, which has been shown to predominately use the late cleavage pathway, shows no difference in infectious virus production when grown on either culture type, again suggesting that it is specifically the serine protease-dependent entry pathway that differs between the culture conditions.

Due to the proprietary nature of commercially available media types, we are unable to determine the factors that drive the differences observed between media types. However, RNA-sequencing of undifferentiated cultures just before ALI shows large differences between cultures grown in either media type, and pathway analysis suggests the growth media is leading to epigenetic differences that persist throughout the differentiation process (supp Fig 4). Additionally, the two different media-grown cultures become more similar over the course of differentiation, likely due to identical genetic background and environment, however the trajectories taken to arrive at the final differentiated culture differ. During differentiation, Ex plus grown cultures upregulate more pathways related to cilia formation and pathways associated with abnormal pulmonary conditions than Airway grown cultures (supp Fig 5A). In contrast, Airway grown cultures upregulate more adhesion related pathways (supp Fig 5B). Additionally, airway-grown cultures specifically downregulate more COVID-19 related pathways during differentiation. Taken together, these data suggest that the epigenetic remodeling both before and after differentiation is impacting the resulting final culture. Future work should investigate the mechanisms of these epigenetic changes to identify factors that may be driving cell fate determination and modulating expression of surface factors.

In conclusion, in this study we show that expansion media influences differentiation patterns and final culture characteristics of airway epithelial cells. We show that this has important implications for SCV2, but not IAV, replication success and is likely due to differences in serine protease cofactor expression. When using these airway culture models for virus studies, and especially therapeutic development, great care should be taken to control for known factors that can influence conclusions-cell type proportion, expression of key proteins, etc ^9,44–46^. However, when working with novel viruses where not much is known, collaboration and independent validation is key to identify confounding variables in these studies and to gain high confidence in conclusions and public health recommendations. While differentiated airway epithelial cell cultures are excellent surrogates for studying the respiratory tract, it is also important to remember that the model has limitations and does not perfect recapitulate the heterogeneity of the respiratory tract ^12^. Therefore, future studies and optimizations will no doubt continue to refine this tool.

## Author Contributions

Conceptualization, A.P. and J.D.R.; methodology, A.P. and J.D.R.; acquisition of data, J.D.R, J.L.W., E.A., A.C., M.L., and W.Z.; formal analysis, J.D.R., J.L.W. and E.A.; resources, A.P.; data curation, J.D.R.; writing—original draft preparation, J.D.R., E.A. and A.P.; writing—review and editing, J.D.R., E.A. and A.P.; visualization, J.D.R.; supervision, A.P.; funding acquisition, A.P. All authors have read and agreed to the published version of the manuscript.

## Acknowledgments

We thank the members of the Nicole Baumgarth, Kimberly Davis, Sabra Klein and Andrew Pekosz laboratories for insightful comments and discussion pertaining to this manuscript. This work was supported by T32GM007814-37 (JR), T32AI007417 (JR), the Johns Hopkins Centers of Excellence for Influenza Research and Surveillance (NIAID N272201400007C), the Johns Hopkins Centers of Excellence in Influenza Research and Response (NIAID N7593021C00045) and the Richard Eliasberg Family Foundation (AP). We thank the Bloomberg Flow Cytometry and Immunology Core for use of the MSD instrument. We also thank Anne Jedlicka and Amanda Dziedzic of the Johns Hopkins Bloomberg School of Public Health Genomic Analysis and Sequencing Core Facility for their help with preparing and sequencing samples.

## Supplement

**Supp Fig 1:**
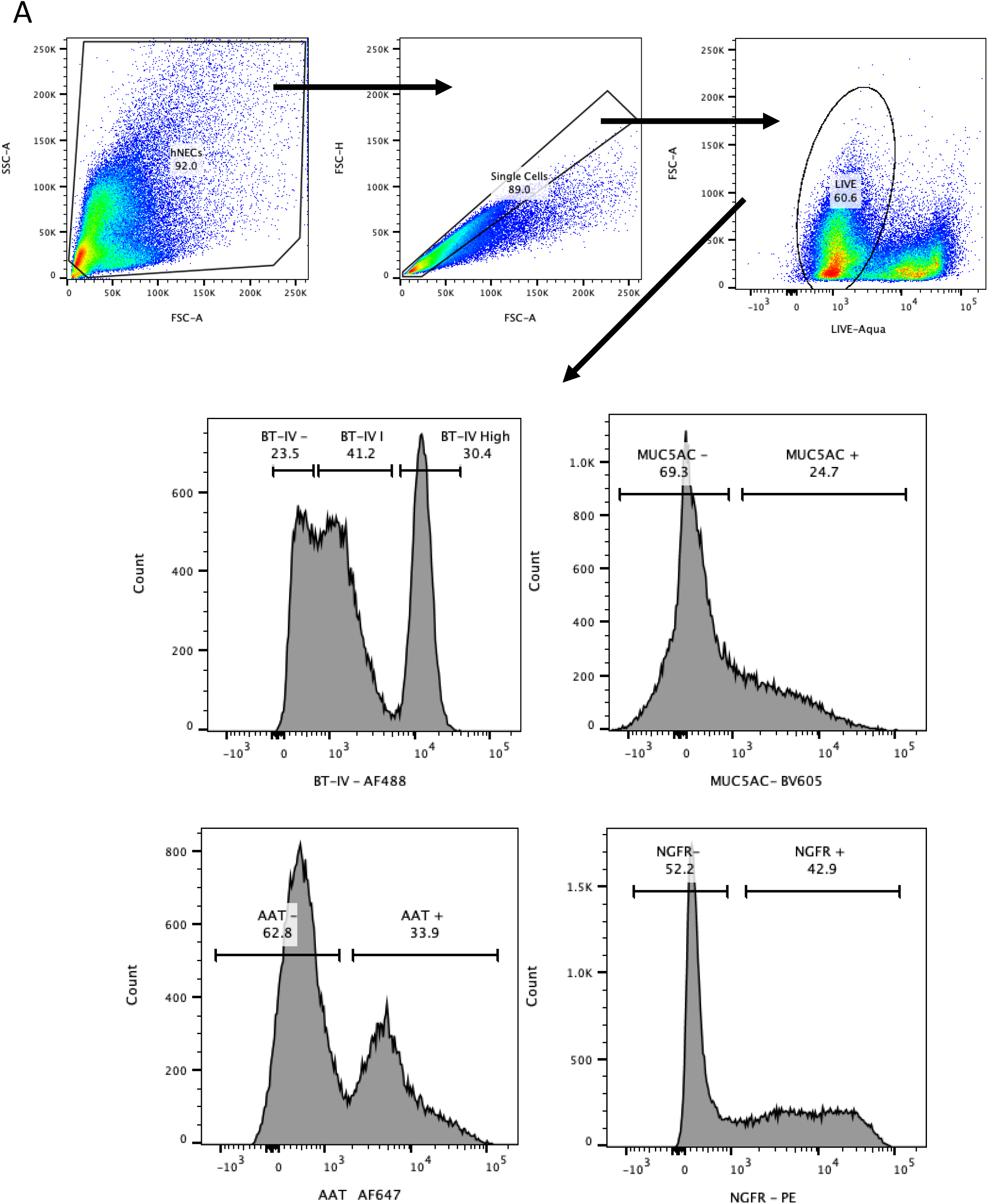
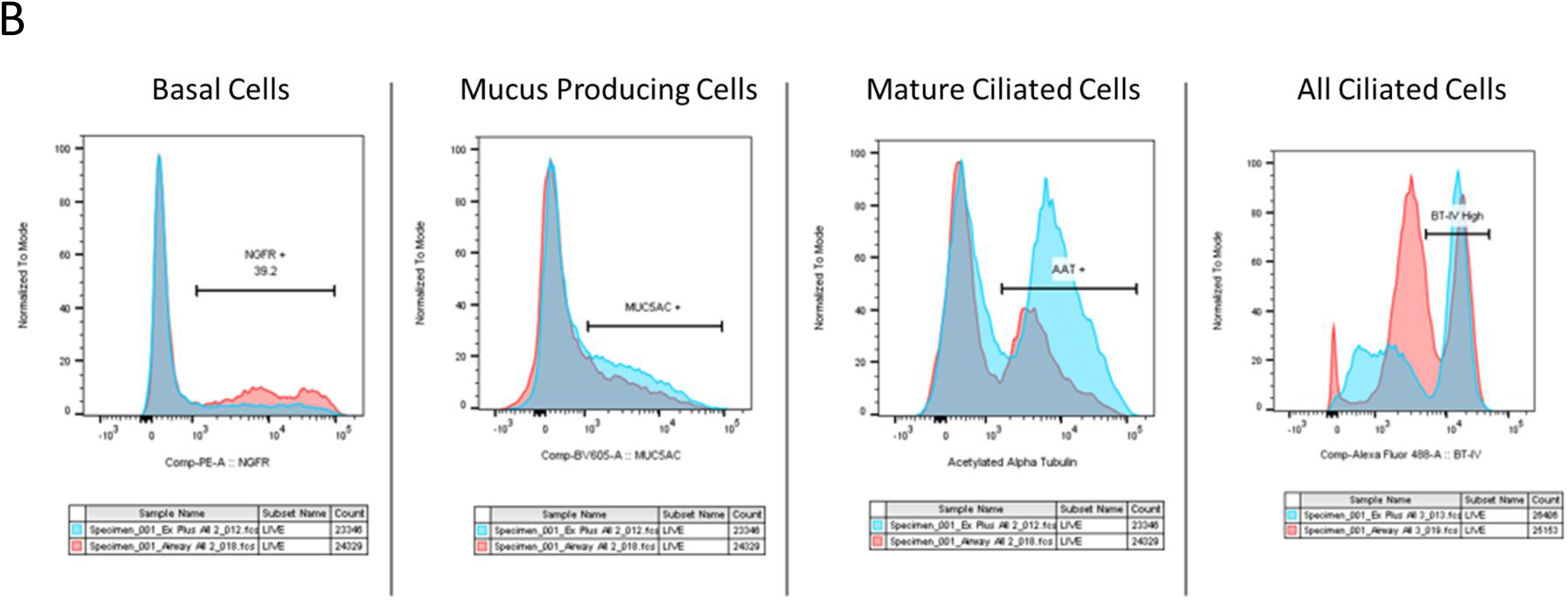
Flow cytometry gating strategy and histogram of shifts in cell type proportion. Ex Plus or Airway cultures were collected for flow cytometry as described. Gating strategy is as shown, first gating on single cells then live cells then presence of cell type markers (A). Proportions of cell types were calculated using histograms of expression (B).

**Supp Fig 2:**
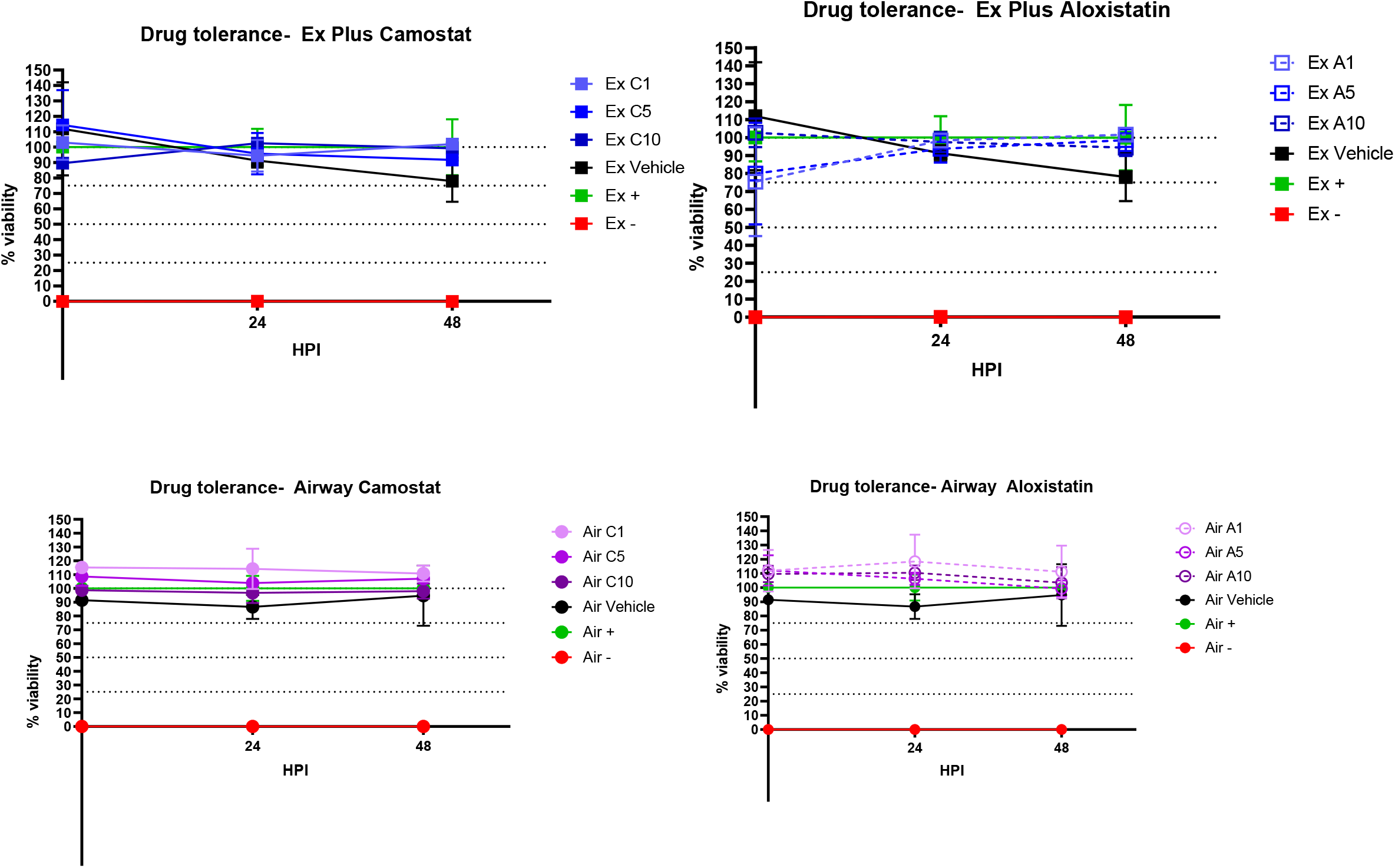
Cell viability after treatment with Camostat or Aloxistatin. Ex Plus (A,C) or Airway(B,D) cultures were pretreated with the indicated concentration of each drug for 24 hours and then viability was measured every subsequent 24 hours until 72 hours had passed. Viability was determined by alamarBlue and based off of a media only and untreated well. Data are pooled from 3 wells per condition and the experiment was repeated once.

**Supp Fig 3:**
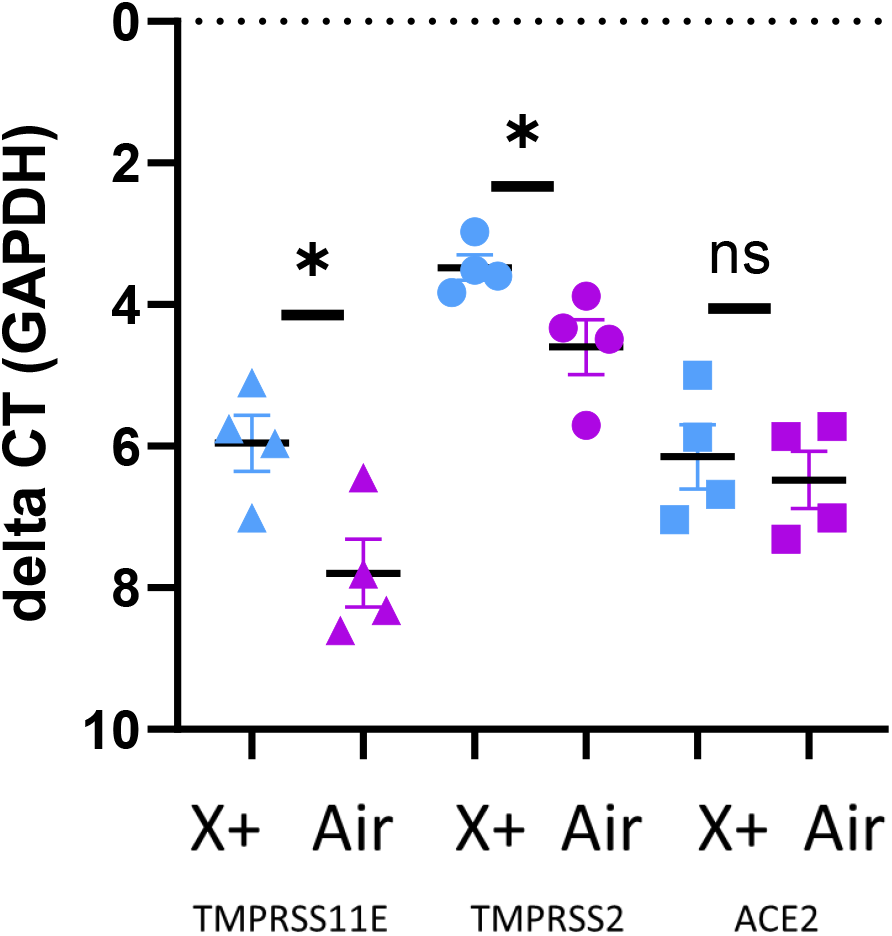
qPCR validation of cofactor and receptor expression in either Airway or Ex Plus grown cultures. Fully differentiated cultures were collected in Trizol and expression of indicated genes was determined using qPCR. N=4 wells from one of two replicate experiments is shown. *P <0.05 (One way ANOVA with Tukey’s posttest)

**Supp Figure 4.**
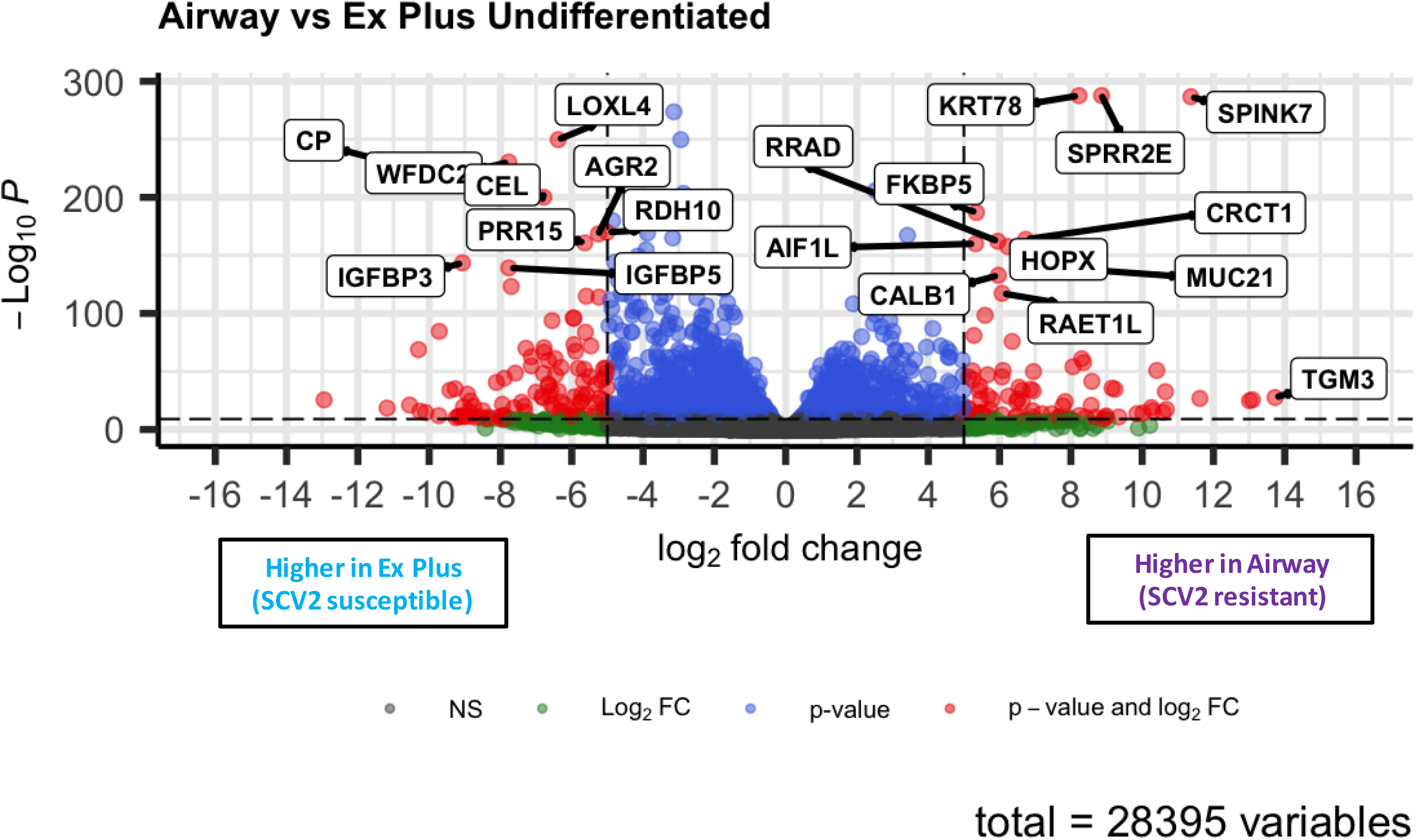
Differentially expressed genes between undifferentiated Ex Plus and Airway expanded cultures. Undifferentiated cultures were collected for bulk RNA-sequencing at day 10-12 when the TEER reading was above 250Ω and transwell was confluent by eye. Data were pooled from three replicate wells. Log 2 fold change indicates the mean expression for a gene. Each dot represents one gene. Black dots indicate no significantly differential expression between Airway and Ex Plus expanded cultures. Blue dots indicate an adjusted p value <10e-10. Green dots indicate an absolute log 2 fold change higher than 5. Red dots indicate both a significant p value and log 2 fold change.

**Supp Figure 5.**
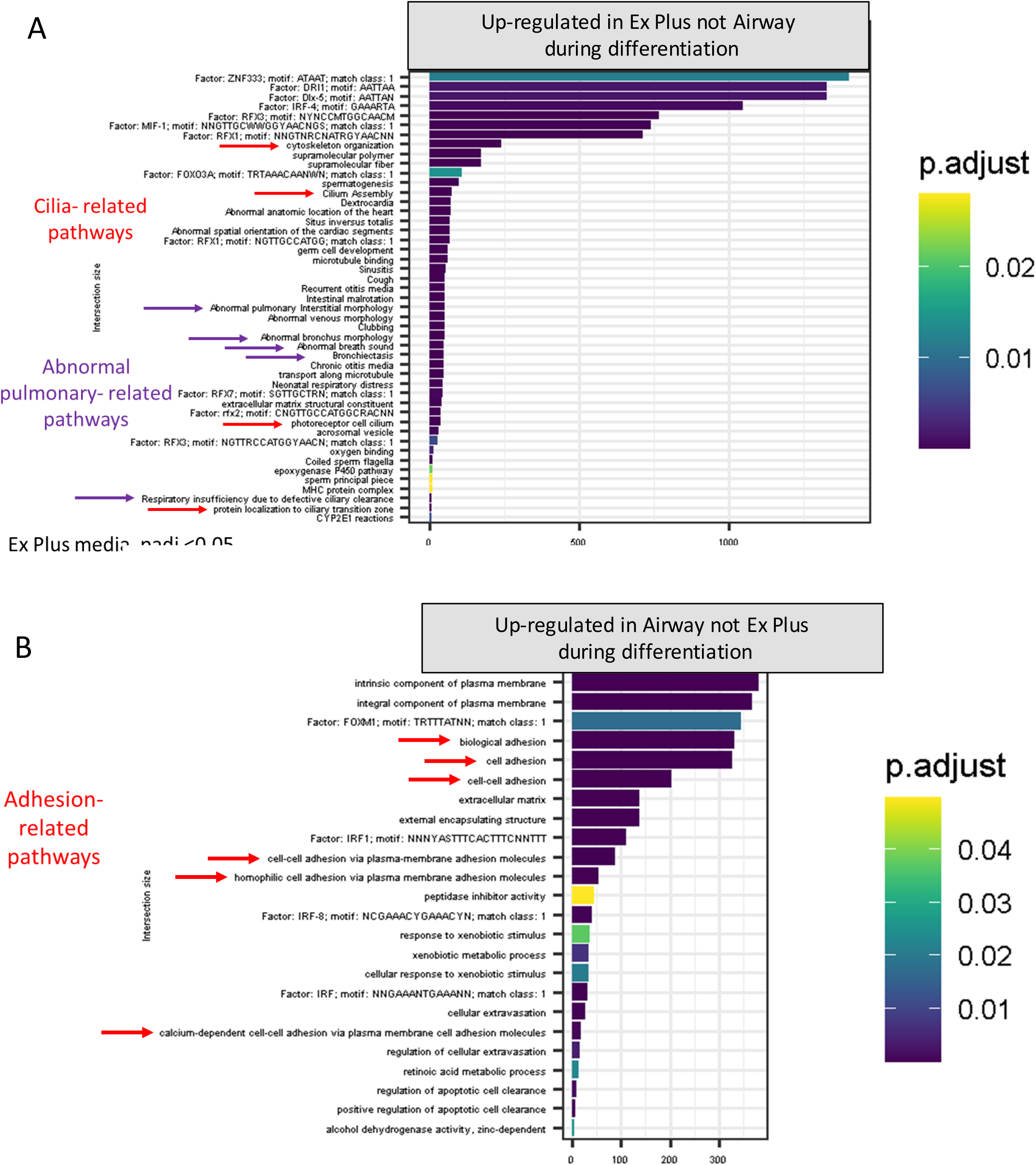
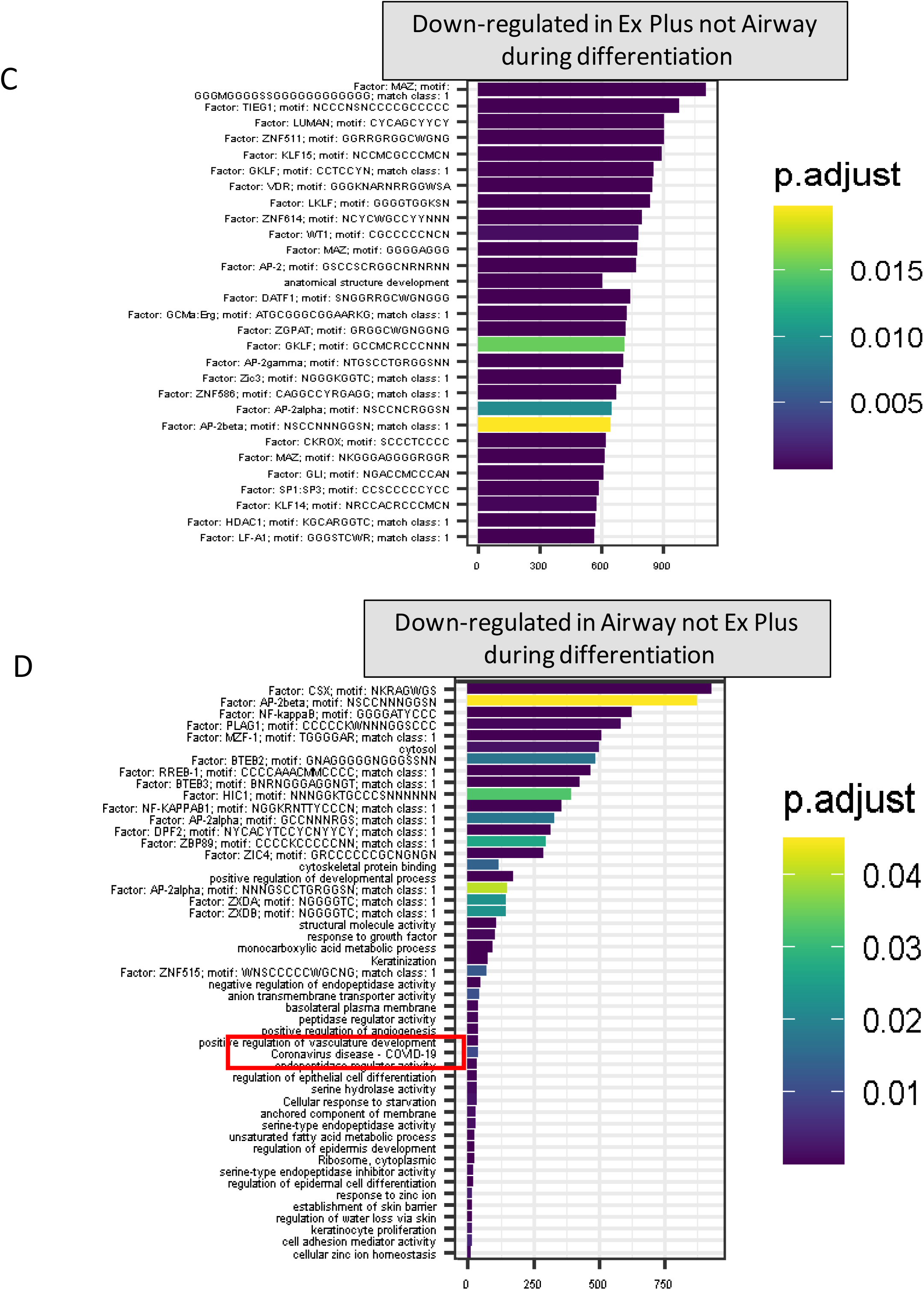
Pathways differentially regulated during differentiation within each culture condition. Undifferentiated cultures were collected for bulk RNA-sequencing at day 10-12 when the TEER reading was above 250 Ωcm and transwell was confluent by eye. Data were pooled from three replicate wells. Significantly differentially expressed genes between undifferentiated and differentiated samples within each expansion media group (p adj <0.05) were used for pathway analysis. Enrichmed pathways were compared between Airway and Ex Plus and pathways unique to each were determined in each direction-up regulated during differentiation (A and B) or down regulated during differentiation (C and D).

